# Targeted deletions in human mitochondrial DNA engineered by Type V CRISPR-Cas12a system

**DOI:** 10.1101/2024.10.20.619292

**Authors:** Natalia Nikitchina, Anne-Marie Heckel, Nikita Shebanov, Ilya Mazunin, Ivan Tarassov, Nina Entelis

**Affiliations:** UMR7156 – Molecular Genetics, Genomics, Microbiology, CNRS/University of Strasbourg, Strasbourg, 67000, France; Department of Biology and Genetics, Petrovsky Medical University, Moscow, 119435, Russia

## Abstract

Mutations in mitochondrial DNA (mtDNA) contribute to various neuromuscular diseases, with severity depending on heteroplasmy level when mutant and wild-type mtDNA coexist within the same cell. Developing methods to model mtDNA dysfunction is crucial for experimental therapies. Here, we adapted the Type V CRISPR-AsCas12a system, which recognizes AT-rich PAM sequences, for targeted editing of human mtDNA. We show that AsCas12a effector, fused with a mitochondrial targeting sequence (MTS) from *Neurospora crassa* ATPase subunit 9, is efficiently addressed into human mitochondria and induces specific mtDNA cleavage in human cells. As a proof-of-concept, we demonstrate that AsCas12a, complexed with two crRNAs targeting distant regions of human mtDNA, introduces specific deletions in mtDNA. For the first time, we provide experimental data proving that a CRISPR system can be used not only for mtDNA degradation but also for precise mtDNA manipulation, offering a potential therapeutic avenue to address mitochondrial disorders.

## Introduction

Mitochondria are double-membrane organelles present in nearly all eukaryotic cells [1]. While traditionally known as the “powerhouse of the cell” for their role in energy production, mitochondria are now recognized as key signaling organelles involved in calcium homeostasis, apoptosis, heme and iron-sulfur cluster synthesis, and even immune regulation [2, 3]. Mitochondria contain their own mitochondrial DNA (mtDNA), from which much of ancestral proteobacterial genome has either been lost or transferred to the nuclear genome over evolutionary time [4]. Human mtDNA represents a 16.5 kb circular DNA encoding 13 subunits of four out of five OXPHOS complexes, 22 tRNAs and 2 ribosomal RNAs [5].

Despite similarities to nuclear DNA (nDNA) in terms of susceptibility to damage, mtDNA repair mechanisms are notably less efficient [6, 7]. This inefficiency is partly compensated by the presence of multiple copies of mtDNA within each cell, which buffers against the negative effect of mutations. Unlike nDNA, where repair is prioritized, damaged mtDNA containing abasic sites [8] or double-stranded breaks (DSBs) [9] is more often rapidly degraded rather than repaired.

Mutations in mtDNA are linked to a variety of neurodegenerative and neuromuscular disorders, as well as phenotypes like infertility, reduced lifespan, heart disease, diabetes and early-onset Alzheimer’s disease [10]. Over 300 distinct mtDNA pathogenic mutations have been identified, contributing to various diseases [11], affecting approximately 1 in 5000 individuals [12]. Disease severity is closely tied to the level of heteroplasmy, the state where both mutant and wild-type mtDNA (WT mtDNA) co-exist within the same cell [11]. Currently, there are no effective treatments for these pathologies [13].

Due to the lack of treatments, extensive research is focused on developing new tools for genetic manipulation of mtDNA. Nowadays, the only approaches proven to efficiently reduce mutant mtDNA levels involve the introduction of DSBs, leading to selective degradation of damaged mtDNA. Among these approaches, mitochondria-targeted zinc-finger nucleases (mitoZFNs) and transcription activator-like effector nucleases (mitoTALENs) have demonstrated efficacy in reducing mutant mtDNA levels [14, 15]. Despite their potential, mitoZFNs and mitoTALENs face challenges, such as the complex design of DNA-binding domains for diverse mtDNA sequences and the size constraints of existing viral delivery systems [16]. Additionally, such systems are not applicable for homoplasmic pathogenic mtDNA mutations, where all copies of mtDNA are mutated.

Despite its clinical relevance, the mechanisms of mtDNA repair after DSBs remain poorly understood. While nDNA repair pathways, such as Homology-Directed Repair (HDR) and Non-Homologous End Joining (NHEJ), are well characterized, mtDNA with DSBs, as already mentioned, is typically degraded rather than repaired [9, 17]. Although some evidence from *in vitro* and *in vivo* studies suggests that HDR, NHEJ, and Microhomology-Mediated End Joining (MMEJ), an alternative repair pathway, which uses small homologous regions to facilitate rejoining, may occur in mitochondria [18–20], their roles and mechanisms remain unclear.

Studies using mitochondrially targeted restriction endonuclease *Pst*I, which cleaves at two sites in human mtDNA, showed that sticky ends can re-ligate, thus forming deletions instead of inducing degradation [21, 22]. This phenomenon suggests that introducing DSBs could be a feasible approach for generating pre-designed deletions in mtDNA. So far, these studies have not elucidated the factors governing such deletions.

The emergence of a multitude of CRISPR-Cas systems has provided a powerful tool for nuclear genome editing [23, 24]. However, their application to mtDNA editing remains a challenge due to the unique properties of mitochondria. A significant hurdle is the absence of a known system for importing guide RNAs (gRNAs) into mitochondria, a key requirement for conventional CRISPR systems like Cas9, which rely on RNA-guided recognition of target sequences [15].

Despite these challenges, recent studies have demonstrated that CRISPR-Cas9 systems, when targeted to mitochondria, can generate DSBs that result in a detectable reduction in mtDNA copy number [25–29]. This reduction in mtDNA copy number following CRISPR-induced DSBs has shown promise as a method to selectively deplete mutated mtDNA, offering potential therapeutic benefits. However, without definitive proof of targeted mutagenesis, it remains uncertain whether CRISPR systems can be effectively used for mtDNA editing.

To address these gaps and further explore the potential of CRISPR systems in mtDNA editing, the current study focuses on employing the Type V CRISPR-AsCas12a system in human cultured cells. Unlike Cas9, Cas12a generates DSBs with staggered cuts, producing sticky ends that are expected to facilitate the re-ligation of intervening DNA sequences, resulting in the formation of deletion [30, 31]. By using two crRNAs designed to target distant regions of mtDNA, we aimed to generate deletions mimicking those observed in mitochondrial disorders. Additionally, since mtDNA molecules bearing DSBs undergo rapid degradation, re-ligation of mtDNA increases its stability and thus facilitates detection of the edited molecules. This strategy also allows us to bypass the RNA import challenge associated with gRNA-dependent systems, as AsCas12a eliminates the need for long gRNAs and instead operates with crRNAs, which are much shorter (40–44 nucleotides) [30]. Indeed, the shorter length of crRNAs makes them a more suitable substrate for mitochondrial import compared to gRNAs, as demonstrated in our recent study, where AsCas12a crRNAs were found to be imported into the mitochondrial matrix without the need for additional mitochondrial targeting signals [32]. The obtained data suggest that CRISPR-AsCas12a system can be applicable to induce deletions at targeted sites, demonstrating its potential as a precise editing tool for human mtDNA.

## Results

### Targeting AsCas12a nuclease into human mitochondria

Proteins can be imported into mitochondria by adding mitochondrial targeting peptides [33]. To target AsCas12a into mitochondrial matrix, we attached various mitochondrial targeting signals (MTSs) to the N-terminus of the nuclease (**Fig. 1a**). Along with one of the most widely used MTSs of cytochrome c oxidase subunit 8 (COX8A), we aimed to compare MTSs derived from various sources. These included MTS of *Neurospora crassa* ATPase subunit 9 (Su9) and MTS of the mammalian autophagy-related cysteine peptidase 4D (ATG4D) [34] and MTS of superoxide dismutase 2 (SOD2) [35] (**Supplementary Table 1**).

**Figure 1.**
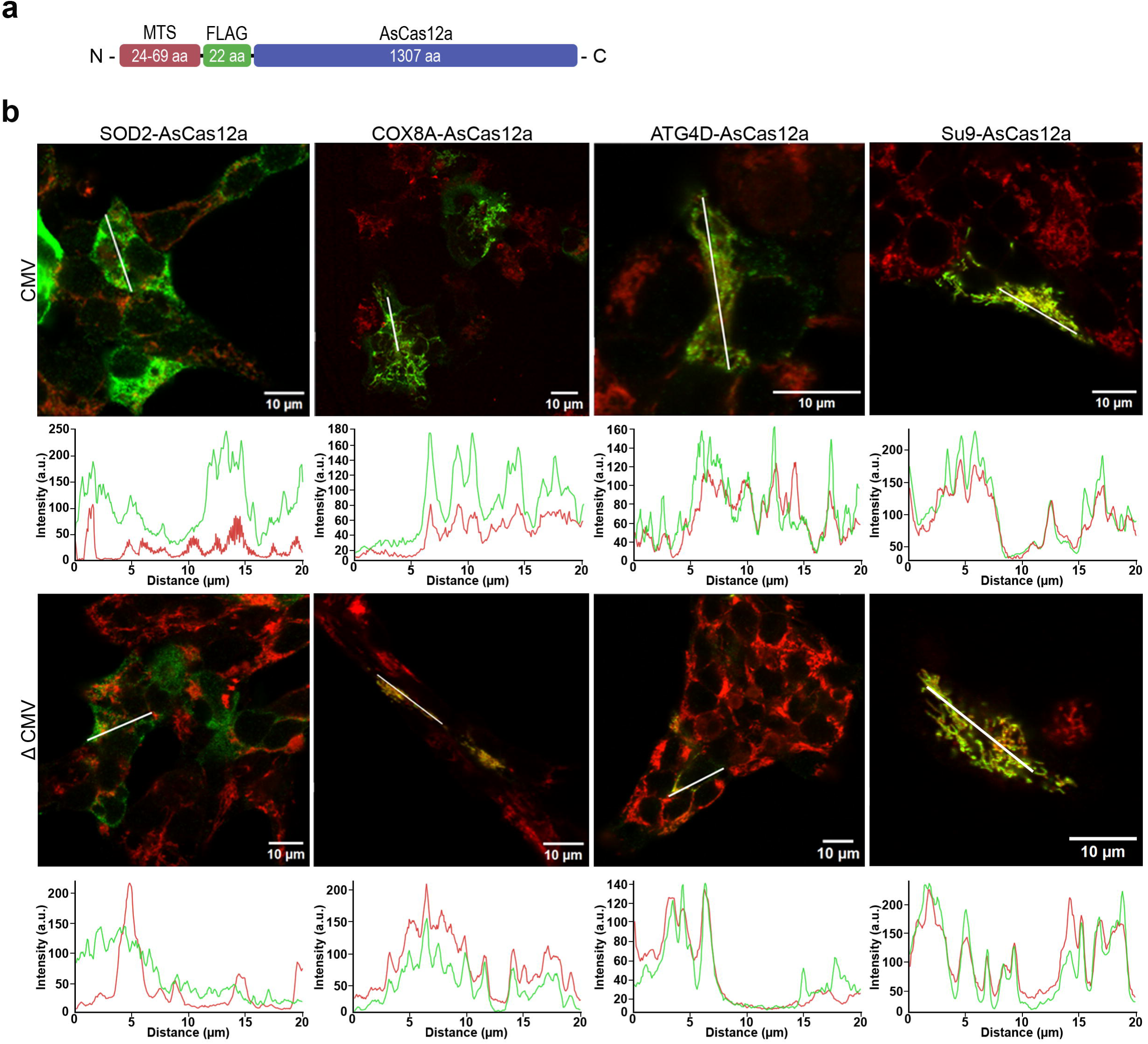
Mitochondrial localization of AsCas12a bearing various MTSs. **a** Scheme representing the construct used. Mitochondrial-targeting signal (MTS) is shown in red, 3xFLAG-tag in green, and the AsCas12a nuclease gene in blue. The amino acid length (aa) is specified within the blocks. **b** Immunofluorescent staining of T-REx-293 cells 24 h after transient transfection with the plasmids bearing AsCas12a fused with various MTSs, as indicated above the panels. AsCas12a fused to different MTSs was expressed under the control of either the full-sized (upper panels) or weakened CMV promoter (lower panels). The 3xFLAG-tagged nuclease was stained with FLAG-tag/Alexa Fluor 488 antibodies (green signal), and mitochondria were stained with TOMM20/Alexa Fluor 647 antibodies (red signal). Images of merged fluorescent signals are represented. To assess mitochondrial colocalization of AsCas12a, a semi-quantitative analysis of the images was performed by measuring the fluorescence profiles of both signals along a 20 *μ*m line (colored in white) (profiles shown under each panel). Scale bars: 10 *μ*m; a.u. -arbitrary units.

To assess the mitochondrial import potential of AsCas12a with the abovementioned MTSs, we analyzed protein properties such as the predicted pI, charge, and grand average of hydropathy (GRAVY) for the amino acid sequences of the MTSs alone and when fused to AsCas12a (**Supplementary Table 2**). All the AsCas12a fusions with various MTSs displayed comparable negative GRAVY values, suggesting that the resulting proteins are expected to possess hydrophilic properties, enhancing their solubility [36]. Regarding the charge, SOD2 MTS appeared less favorable for fusion with AsCas12a due to its relatively small positive charge, and even a negative charge in the full protein context. COX8A MTS emerged as a promising candidate; ATG4D and Su9 MTSs were anticipated to be the most effective for AsCas12a delivery owing to their notably high positive charge. These predictions should be considered with caution, as they are based on analyzing the amino acid sequence, and their accuracy can vary depending on protein folding.

To experimentally test these predictions, we created eight plasmids, each carrying AsCas12a fused with one of four different MTSs (**Fig. 1a**, **Supplementary Table 1**), under the regulation of either the full-sized or shortened CMV promoter region. The AsCas12a expression from a weakened (Δ5) CMV promoter [37] was applied to address potential protein aggregation in the cytosol due to overexpression [38]. The decrease in the nuclease expression under the control of the weaker promoter was confirmed by Western blot analysis (**Supplementary** Fig. 1). Next, T-REx-293 cells transfected with these plasmids were examined by immunocytochemistry (**Fig. 1b**). As predicted, AsCas12a fused with SOD2 MTS exhibited the poorest colocalization profile regardless of the expression level. The proteins bearing COX8A and ATG4D MTSs demonstrated improved mitochondrial localization, though they were still partially localized in the cytosol. Notably, reducing the expression of AsCas12a versions improved their mitochondrial localization (**Fig. 1b**). Finally, AsCas12a fused with Su9 MTS demonstrated the best mitochondrial colocalization profile.

### Developing a human cell line expressing mitochondrial-targeted AsCas12a

Based on the above results, we developed a stable cell line expressing the nuclease fused with Su9 MTS and being expressed in a tetracycline-inducible manner (**Fig. 1b**); 100 ng/mL of tetracycline was determined as the optimal concentration for further experiments (**Supplementary** Fig. 2a). We then assessed the stability of Su9-AsCas12a in cells over time by analyzing total cell lysates at several time points within a week after the nuclease expression activation (**Supplementary** Fig. 2b). The results revealed that AsCas12a was still present in cell lysates even on the sixth day after induction, although the amount decreased over time.

To confirm mitochondrial localization of Su9-AsCas12a in the stable cell line, we performed immunofluorescent staining after the nuclease expression activation. The results confirmed our initial observation on the mitochondrial localization of Su9-AsCas12a (**Fig. 2a**).

**Figure 2.**
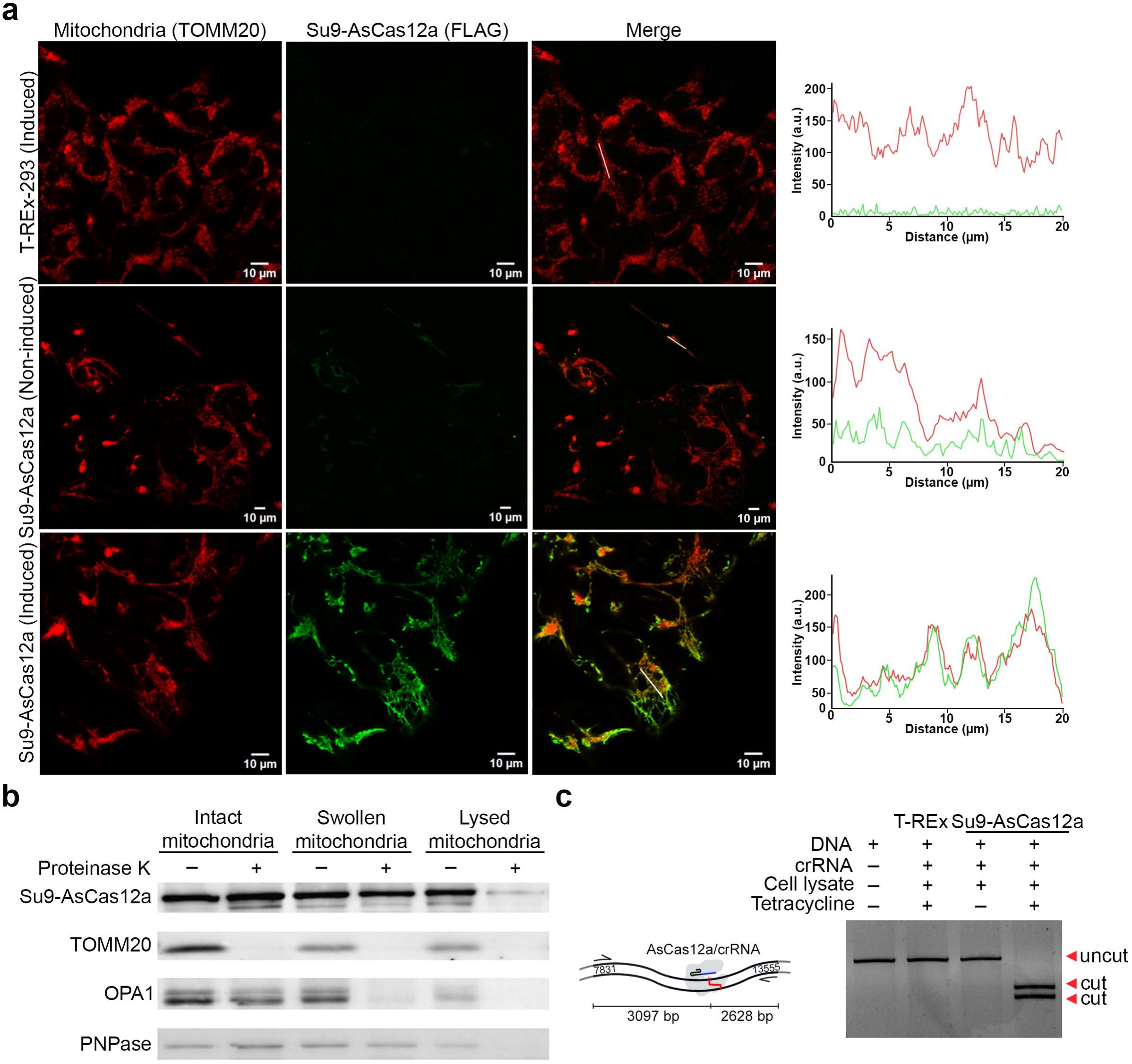
Su9-AsCas12a expressed in the stable cell line is localized in mitochondrial matrix and possesses its specific DNA cleavage activity. **a** Immunofluorescent staining of the T-REx-293-Su9-AsCas12a monoclonal stable cell line 24 h after the nuclease expression activation (induced). Non-induced stable cell line and induced T-REx-293 cells were used as controls. The nuclease was labeled with FLAG-tag/Alexa Fluor 555 antibodies (green signal), while mitochondria were labeled with TOMM20/Alexa Fluor 647 antibodies (red signal). Colocalization of the fluorescent signals were assessed as in Fig. 1b. **b** Submitochondrial localization of Su9-AsCas12a in the created stable cell line. Intact mitochondria, swollen mitochondria (mitoplasts), and lysed mitochondria were treated or not with proteinase K (as indicated above the panel), and the residual proteins were analyzed by Western blotting. The antibodies used are shown on the left: FLAG-tag (epitope tag fused to AsCas12a); PNPase (intermembrane space and matrix-localized mitoribosomal protein); TOMM20 (mitochondrial outer-membrane protein facing the cytosol); OPA1 (inner membrane protein facing the intermembrane space). Nuclease expression was activated by tetracycline. **c** On the left, schematic representation of a target DNA (PCR fragment of human mtDNA, nucleotide numbers are indicated) containing the AsCas12a PAM sequence; cleavage site is marked with red line. The sizes of the fragments corresponding to the cleaved DNA template are given below the scheme. On the right, target DNA cleavage in cell lysates of the T-REx-293-Su9-AsCas12a stable cell line (‘Su9-AsCas12a’) induced with tetracycline. DNA template (uncut) and its fragments (cut) were separated in an agarose gel. DNA template and crRNA-1 targeting MT-ND4 were added directly to the reaction (indicated above the gel). Lysate from the induced T-REx-293 cell line (‘T-REx’) served as a negative control.

To complement the microscopy data, we performed submitochondrial fractionation of proteins (**Fig. 2b**). Notably, Su9-AsCas12a remained proteinase-resistant in intact and swollen mitochondria (mitoplasts), degrading only when mitochondria were fully lysed. This behavior mirrored that of the matrix-localized protein PNPase [39], thus confirming localization of Su9-AsCas12a in the mitochondrial matrix. In contrast, the outer membrane protein TOMM20, which faces the cytosol [40], was digested in all treated samples, whereas the inner membrane protein OPA1, facing intermembrane space [41], disappeared in treated mitoplasts and lysed mitochondria (**Fig. 2b**).

Next, we evaluated the Su9-AsCas12a nuclease activity in the stable cell line by site-specific cleavage of a DNA template in cell lysate (**Fig. 2c**). The presence of cleavage products of the expected size indicated that Su9-AsCas12a expressed in the stable cell line retained specific DNA cleavage activity.

### Impact of the Su9-AsCas12a expression on human mitochondria

After demonstrating mitochondrial import and activity of Su9-AsCas12a in the stable cell line, we sought to ensure that activating the nuclease expression did not alter the cellular phenotype.

AsCas12a and its orthologs are known to possess trans-cleavage activity, meaning they can degrade single-stranded DNA molecules in a non-specific manner upon binding to the target DNA [42]. Considering the presence of R-loops in human mtDNA [43], the import of small non-coding RNAs such as miRNAs and siRNAs into mitochondria [44], and our recent discovery that AsCas12a crRNA can be divided into scaffold and spacer parts without affecting its cleavage activity [45], one could hypothesize that targeting AsCas12a to human mitochondria might induce non-specific hydrolysis of mtDNA, leading to its depletion, decrease of mitochondrial respiration, and subsequently triggering mitophagy [46].

Based on the fluorescent microscopy data (**Fig. 2a**), we observed that mitochondria form a network of tube-like structures in the stable cell line, similar to those in the control cells, and this network was not affected by the nuclease expression. To further investigate the mitochondrial phenotype and determine whether Su9-AsCas12a expression in the stable cell line negatively impacts mitochondrial respiration, we performed oxygen consumption rate (OCR) measurements in both T-REx-293 and T-REx-293-Su9-AsCas12a cell lines after treatment with tetracycline. Compared to the control cells, the Su9-AsCas12a stable cell line exhibited lower oxygen consumption rates (**Fig. 3a**). Furthermore, both the basal and uncoupled respiration rates in the stable cell line were identical, indicating that these cells were respiring at their maximal capacity [47].

**Figure 3.**
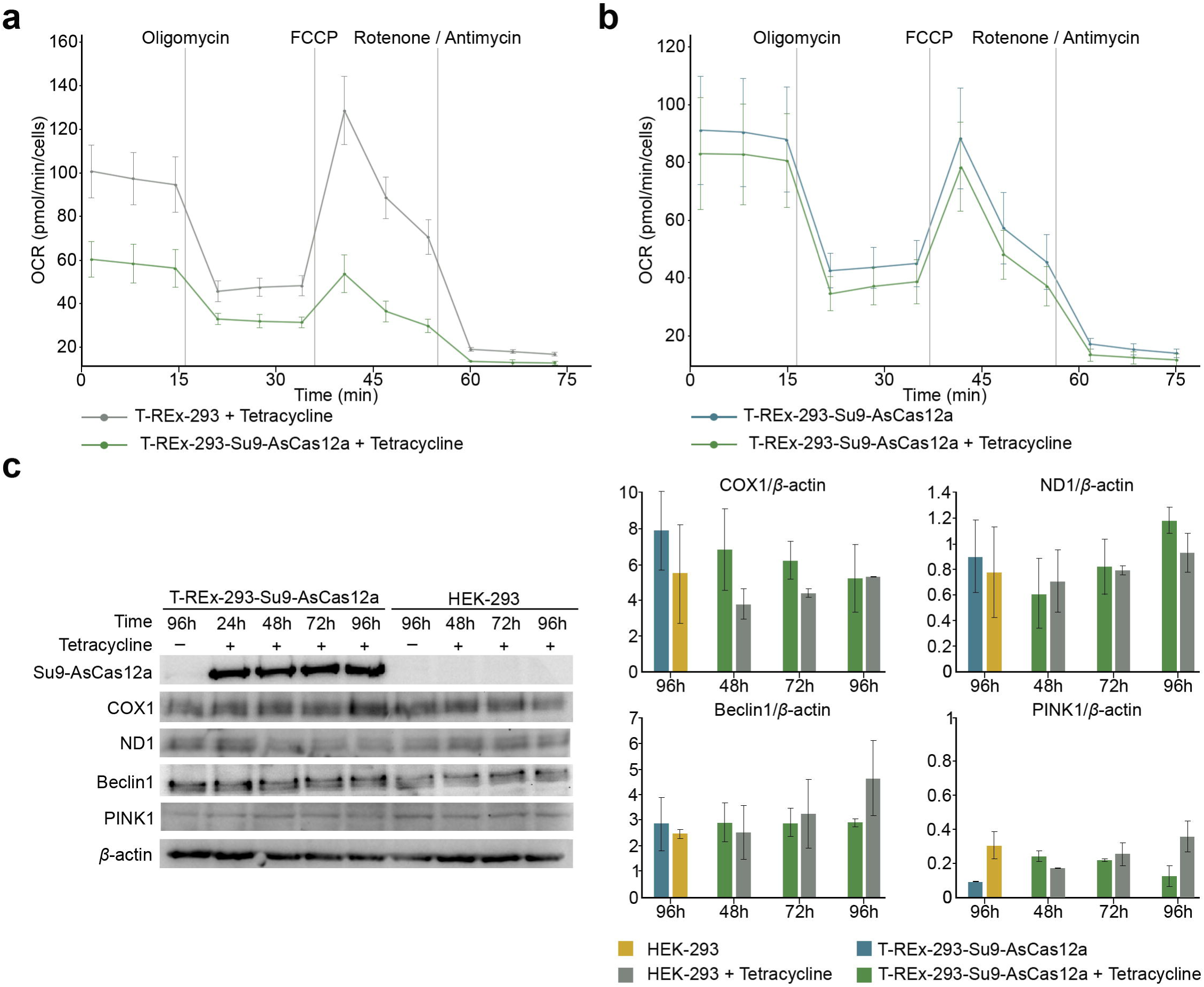
Impact of Su9-AsCas12a expression on mitochondrial respiration, protein expression and mitophagy. **a** Oxygen consumption in T-REx-293 cells (grey line) and the T-REx-Su9-AsCas12a cells (green line) 48 h after the tetracycline treatment. Values and error bars reflect the mean ± s.d. of n = 3 independent biological replicates. **b** Same as **a**, but oxygen consumption in the T-REx-Su9-AsCas12a cells either treated (green line) or not (blue line) with tetracycline. **c** Western blot analysis of total cell lysates of the T-Rex-Su9-AsCas12a stable cell line, cultivated during 24 to 96 h after tetracycline induction (as indicated above the panels). Antibodies (indicated at the left): COX1 and NFD1 (mitochondrially encoded and localized), markers of mitophagy, Beclin 1 and PINK1. As a control, HEK-293 cells were treated in the same way. Quantifications of the steady-state levels of mitochondrial proteins normalized to *β*-actin reflect the mean ± s.d. of n = 2 independent biological replicates.

To determine if this decrease in the respiration rate in the stable cell line is due to the nuclease expression, we compared OCR in the T-REx-293-Su9-AsCas12a cell line with and without the nuclease expression (**Fig. 3b**). The results showed that there is no link between the lower OCR levels and Su9-AsCas12a expression, indicating that decreased respiration rates in the stable cell line cannot be attributed to Su9-AsCas12a mitochondrial targeting.

We then aimed to investigate whether expression of Su9-AsCas12a and its import into human mitochondria could lead to alterations in mitochondrial translation and/or induce mitophagy. We performed Western blotting using total cell lysates from the Su9-AsCas12a stable cell line, induced with tetracycline, to explore whether the nuclease expression affects mitochondrial proteins COX1 and ND1 (mitochondrially encoded proteins localized in the inner membrane respiratory complexes) and the markers of mitophagy Beclin 1 and PINK1 [48]. The levels of COX1 and ND1 proteins did not change upon expression of the mitochondrial-targeted AsCas12a nuclease (**Fig. 3c**). Similarly, no differences were observed in the levels of PINK1 and Beclin 1 (**Fig. 3c**). These findings suggest that when Su9-AsCas12a is expressed and targeted to mitochondria, it neither affects mtDNA expression nor induces mitophagy.

### Functional assessment of the mitoAsCas12a system in living cells

The Type V CRISPR-AsCas12a system cleaves DNA at nucleotide positions 17–19 downstream of the PAM on the non-target DNA strand (NTS), as well as at positions 21–23 on the target strand (TS), generating 3–5 nucleotide-long 5ʹ overhangs [49]. To study the possibility of inducing a deletion in human mtDNA using the mitoAsCas12a system [31], we selected two crRNAs which program mtDNA cleavage generating complementary 5’-overhangs capable of ligation, referred to as crRNA-1 and crRNA-2 (**Fig. 4a**, **4b**).

**Figure 4.**
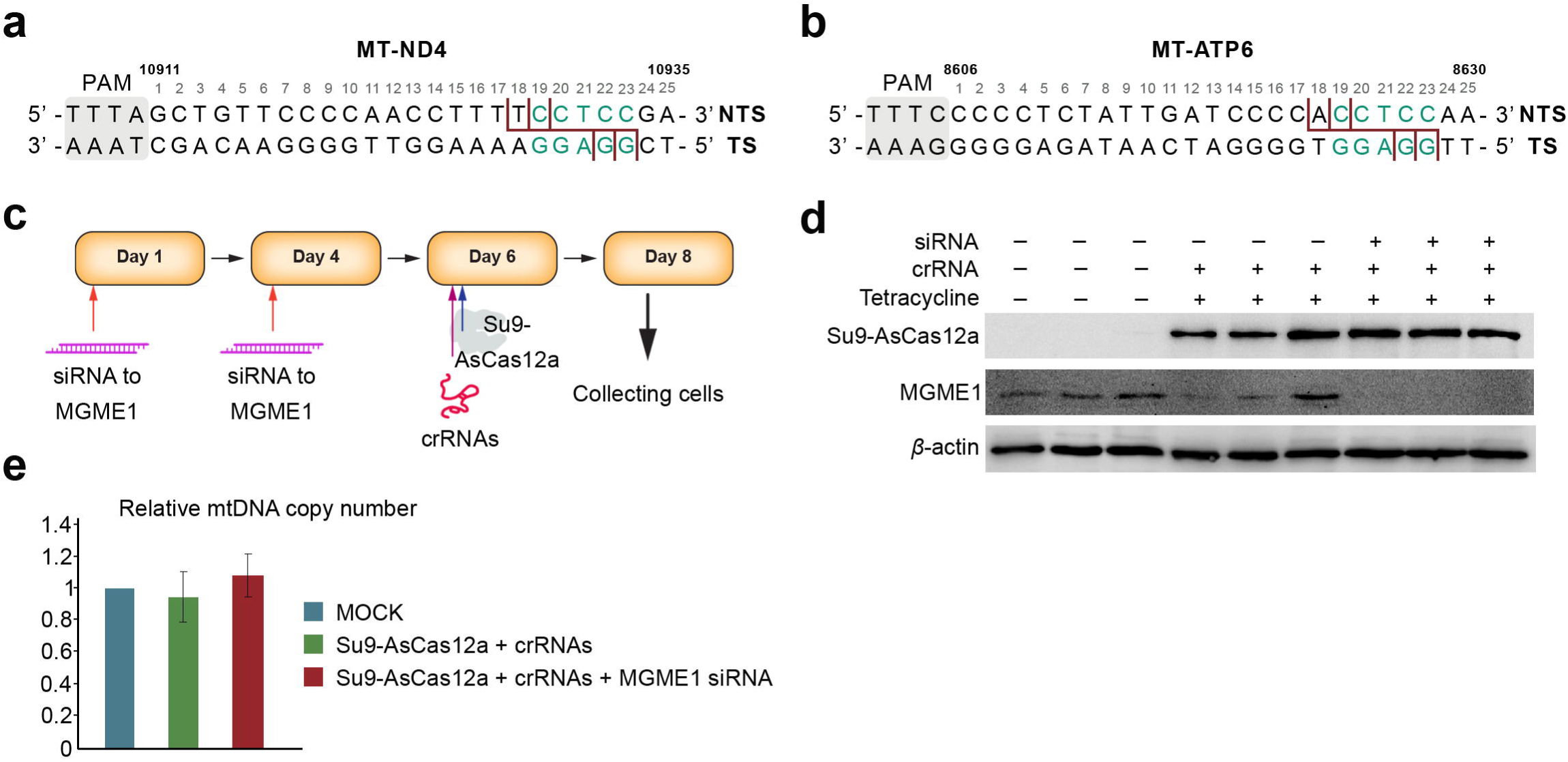
Strategy to induce deletion in human mtDNA. **a, b** Sequences of the sites in human mtDNA targeted by crRNA-1 (**a**), and crRNA-2 (**b**), which were designed to generate complementary ends (corresponding nucleotides are highlighted in green). PAM sequences are shown in grey; NTS, non-target DNA strand; TS, target DNA strand. Possible cleavage sites on both strands are indicated with red lines. The nucleotide coordinates corresponding to the WT human mtDNA are shown above the sequences. **c** Experimental pipeline for the T-REx-Su9-AsCas12a cells transfection to induce a deletion in human mtDNA. **d** Western blot analysis of the cells on Day 8, with or without MGME1 knockdown, in triplicates, as indicated above the panel. Proteins detected are shown at the left. **e** qPCR analysis of relative mtDNA copy number in the T-REx-Su9-AsCas12a stable cell line (‘Su9-AsCas12a’) with or without MGME1 knockdown, Day 8 (shown in red). mtDNA copy number in non-transfected cells (MOCK) is taken as 1. Mean values with standard deviations are presented; n = 3.

The cleavage activity of AsCas12a programmed with these crRNAs was previously demonstrated in cell lysates containing the nuclease [32]. Before testing the system in living cells, and to prevent the degradation of cleaved mtDNA thereby facilitating deletion formation, we treated the stable cell line with an siRNA targeting MGME1, a mitochondrial 5’-3’ exonuclease, which plays a key role in the degradation of linearized mtDNA [9]. Following this treatment, cells were transfected with the two crRNAs (**Fig. 4a, 4b**), and Su9-AsCas12a expression was activated, as shown in **Fig. 4c**. The expression of Su9-AsCas12a and knockdown of MGME1 were confirmed via Western blotting (**Fig. 4d**).

Since MGME1 downregulation can affect not only linearized mtDNA degradation but also mtDNA replication [50], we assessed the mtDNA copy number in cells expressing the mitoCRISPR-Cas12a system. No significant changes in mtDNA copy number were observed regardless of MGME1 knockdown (**Fig. 4e**). This also indicates that the import of the mitoCRISPR-Cas12a system into human mitochondria does not have a mito-toxic effect, consistent with our earlier data (**Fig. 3**).

To detect mtDNA cleavage, we applied a sensitive approach to identify free ends of mtDNA, nested linker-mediated PCR (LM-PCR) (**Fig. 5a**). We detected the expected PCR products corresponding to specific mtDNA cleavage induced by Su9-AsCas12a programmed with crRNA-1 and with crRNA-2 in the stable cell line (**Fig. 5b, 5c**). Noteworthy, we were unable to detect linearized mtDNA molecules in the Su9-AsCas12a cells transfected with crRNA-1 and bearing fully active MGME1, likely due to their rapid degradation (**Fig. 5b**). In the cells transfected with crRNA-2, free ends of mtDNA were detectable in samples not treated with siRNA, though their quantity was strongly reduced compared to samples where MGME1 expression was downregulated (**Fig. 5c**). These data clearly indicate that specific mtDNA cleavage can be induced by the mitoAsCas12a system in cultured cells.

**Figure 5.**
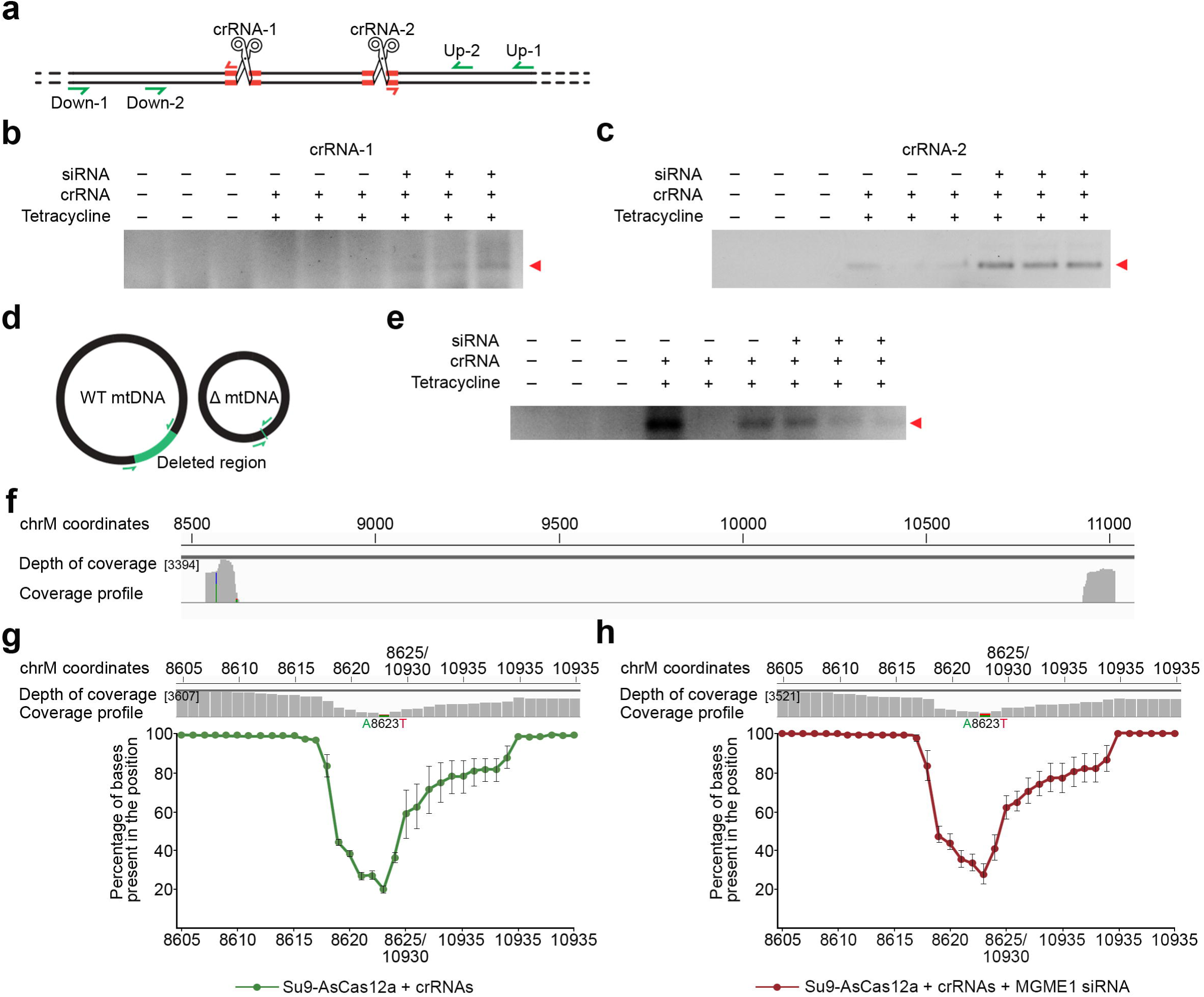
mitoCRISPR system induced specific deletion in mtDNA. **a** Scheme of nested LM-PCR, where mtDNA were ligated with an asymmetric double-stranded linker (shown in red) to identify cleavage sites. Nested LM-PCR involves two rounds of PCR using primers specific to the linker (red arrows) and the mtDNA regions (green arrows). **b** Agarose gel analysis of nested LM-PCR with primers downstream of the crRNA-1 targeting site. Cells treatment is indicated above the gel, samples represent independent biological triplicates. LM-PCR products of the expected size are indicated with red arrow. **c** Similar to (**b**) but using upstream primers to detect cleavage induced by AsCas12a in complex with crRNA-2. **d** Schematic representation of detecting mtDNA molecules bearing a deletion by PCR with primers (green arrows) flanking the deletion (highlighted in green). **e** PCR on total DNA with primers flanking the deletion. Cells were treated as indicated above the gel. Products of the expected size were visualized on an agarose gel (red arrow). **f** Example of the coverage profile for the sample containing Su9-AsCas12a and crRNAs 1 and 2 with MGME1 downregulation, obtained by NGS of PCR amplicons and visualized using IGV. Alignment to the WT human mtDNA. The depth of coverage and chrM coordinates are indicated, with mutations at positions A8566C (40 % of mutation) and A8623T (30 % of mutation) highlighted in color. **g, h** Similar to (**f**), alignment to mtDNA reference bearing the expected 8624/10928 bp deletion for the samples treated with Su9-AsCas12a and crRNAs 1 and 2, either without (**g)** or with (**h**) MGME1 downregulation. Below the coverage profiles, graphs represent the percentage of bases at the nucleotide position of the aligned reads (with values less than 100 % indicating the presence of a deletion at that position). Error bars represent the mean ± s.d. of n = 2 for (**g**) and n = 3 for (**h**) independent biological replicates.

To detect the presence of the expected mtDNA deletion, we performed PCR with primers flanking the deletion boundaries (**Fig. 5d**). A DNA band corresponding to the amplification of mtDNA molecules bearing the deletion was observed in samples treated with both components of the mitoCRISPR system, namely Su9-AsCas12a and crRNAs (**Fig. 5e**). Interestingly, the deletion was also detected in two samples from the biological triplicate without MGME1 downregulation.

Our next objective was to verify the sequence of the obtained amplicons to confirm the formation of the deletion and identify the ligation boundaries. After performing NGS on the amplicons, the resulting reads were aligned to the reference human mtDNA genome (**Fig. 5f**). Upon initial inspection of the coverage profiles, we observed that reads aligned precisely at the primer binding sites located at 8542 and 11015 bp, following by a notable decrease in coverage depth observed at the sites of expected cleavage by AsCas12a in complex with crRNAs 1 and 2 (**Fig. 4a, 4b**; **Fig. 5f**). Additionally, two mismatch mutations were identified: A8566C (MT-ATP6/8) (see also **Discussion section**) and A8623T (MT-ATP6), corresponding to mtDNA nucleotide positions 8623/10928 at the deletion boundaries (discussed below).

To examine the deletion boundaries more closely, the obtained reads were realigned to a reference mtDNA sequence bearing the 8624/10928 bp deletion, predicted in case of successful cleavage of mtDNA by Su9-AsCas12a in complex with crRNAs 1 and 2 and followed by ligation (**Fig. 5g, 5h**). In cells treated with the mitoCRISPR system, we indeed observed the expected deletion, indicating that AsCas12a/crRNAs targeted to human mitochondria were active *ex vivo* (**Fig. 5g**). Additionally, we noticed a drop in coverage in proximity of the crRNAs cleavage sites (positions 8623-8628 and 10928-10933), suggesting that the extremities of mtDNA resulting from the cleavage were slightly degraded prior to ligation. This degradation was more pronounced towards the PAM site of crRNA-2 (**Fig. 4b**). Surprisingly, a similar degradation profile of mtDNA free ends was observed in samples with inhibited MGME1 activity (**Fig. 5h**). Moreover, in both types of samples, as mentioned earlier (**Fig. 5f**), we detected an A-to-T substitution at position 8623 with a frequency of 30 % (**Fig. 5g, 5h**). This could be attributed to variations at the AsCas12a nuclease cleavage site, namely to mtDNA cleavage occurring partially at the position 17 downstream of the PAM site, as illustrated in **Fig. 4a, 4b**. It is important to note that our estimation of the deletion boundaries should be interpreted with caution, as sequencing mtDNA amplicons may introduce a bias toward certain variants. Nevertheless, we demonstrated for the first time that a deletion in human mtDNA can be induced by a CRISPR-Cas system targeted to mitochondria.

To further confirm the ability of the mitochondria-targeted CRISPR-Cas system to induce deletions in mtDNA, we decided to generate a larger deletion of 7 kb (8624/15738 bp) using crRNA-3 targeting MT-CYB in combination with crRNA-2 targeting MT-ATP6, which are also able to create cohesive overhangs (**Supplementary** Fig. 3a**, 3b**). Using the optimized pipeline, we treated the stable cell line expressing Su9-AsCas12a with crRNAs 2 and 3, along with siRNA against MGME1 exonuclease (**Fig. 4c**). First, we verified the cleavage induced by the new crRNA-3 targeting MT-CYB using nested LM-PCR (**Supplementary** Fig. 3c**, 3d**). mtDNA molecules bearing the newly induced deletion were detected and analyzed as described above by PCR with primers flanking the deletion, followed by NGS. Alignment to the WT mtDNA reference sequence confirmed that the drop in coverage corresponded to the cleavage positions induced by the nuclease and crRNAs (**Supplementary** Fig. 3e).

Upon precise analysis of the deletion boundaries, we observed a drop in coverage near the crRNAs 2 and 3 cleavage sites, which could result from the degradation of linearized mtDNA prior free-ends ligation (**Supplementary** Fig. 3f, **3g**). However, unlike our previous observations, no SNPs were detected near the cleavage sites.

In summary, for the first time, using next-generation sequencing, we demonstrated that a deletion in human mtDNA can be induced by a CRISPR-Cas system targeted to mitochondria.

## Discussion

Along with point mutations, deletions in mtDNA are a primary cause of mitochondrial diseases, and they also contribute to tissue aging [51]. Of note, the first identified mtDNA mutation was a large-scale 4,977 bp “common deletion”, which has been linked to various diseases and accounts for approximately 12 % of mitochondrial disorders in adults [52, 53]. Deletions in mtDNA can vary in size from 1.8 to 8 kb and can occur almost anywhere in the mitochondrial genome. However, the precise mechanisms behind deletion formation and dynamics remain unknown [54]. Developing novel methods to model mtDNA dysfunction in cell cultures and living organisms is crucial to advance experimental therapy for the long-term treatment of mitochondrial diseases. To date, only a limited number of murine models of mtDNA diseases have been developed [55, 56], including the ΔmtDNA4696 “mito-mice,” which represent the first and only heteroplasmic mouse model with a pathogenic mtDNA deletion [57].

In recent years, CRISPR-based technologies have emerged as powerful tools for genome editing, including potential applications in mitochondrial gene therapy. However, the lack of crucial evidence of such systems activity raises fundamental concerns regarding the applicability of CRISPR systems for mtDNA manipulation and therapeutic interventions [15]. Mitochondrial gene therapy remains challenging because of the double-membrane barrier and the absence of the clear mechanism of RNA import into human mitochondria [58]. Nevertheless, recent advances in CRISPR technologies have demonstrated the potential to address these challenges. The AsCas12a system, a Type V CRISPR nuclease, has emerged as a promising candidate for targeted genome editing [42]. Unlike Cas9, AsCas12a induces staggered cuts, producing 5’ overhangs that can be exploited for generating specific deletions. Additionally, AsCas12a has slightly smaller protein size and requires shorter crRNAs to be programmed, compared to those of Cas9, which makes this system a more promising candidate for mitochondrial targeting [30, 32].

Here, we aimed to investigate whether AsCas12a, when fused with an MTS, could be successfully imported into mitochondria and used to generate specific mtDNA deletions. To date the only attempt to optimize Cas12a for mtDNA targeting was initiated by **Antón et al. in 2020** [34], in which GFP-fused Cas12a bearing different MTSs were compared, but no significant effect on human mtDNA was observed.

To target AsCas12a to the mitochondrial matrix, we fused the nuclease with various MTSs. Although COX8A and SOD2 MTSs are among the most commonly used for directing proteins into mitochondria, they showed poor mitochondrial localization in transiently transfected cells (**Fig. 1b**). In contrast, Su9 MTS provided the most efficient mitochondrial import, as demonstrated by immunofluorescence staining and Western blotting in both transfected cells and the stable cell line (**Fig. 1b**; **Fig. 2a, 2b**). This result aligns with our prediction that Su9 MTS would function as the ‘strongest’ presequence, given that it is longer, more positively charged, and less hydrophobic (**Supplementary Table 2**). Longer MTSs have been shown to contain segments that enhance binding to TOM receptors, thereby increasing import efficiency [59].

Notably, controlling expression levels using a weakened promoter further improved mitochondrial targeting, reducing cytosolic mislocalization (**Fig. 1b**). These findings underscore the importance of both MTS selection, consistent with previous studies linking protein charge to mitochondrial targeting [33], and expression level control [60] in optimizing the mitochondrial import of targeted proteins.

It has previously been shown that prolonged expression of mitochondria-targeted LbCas12a negatively affects mitochondrial respiration [34]. Thus, we assessed whether Su9-AsCas12a expression also affects mitochondrial functions in our experimental setup and observed that the nuclease expression did not alter mitochondrial network morphology (**Fig. 2a)**, as well as steady-state levels of mitochondrial proteins (COX1 and ND1) and mitophagy markers (Beclin 1 and PINK1) (**Fig. 3c**). Regarding mitochondrial respiration, although the Su9-AsCas12a-expressing cell line exhibited a lower OCR compared to control HEK293T cells (**Fig. 3a**), no difference was observed between non-induced and induced states of the stable cell line (**Fig. 3b**). This clearly indicated that the reduced OCR was not a consequence of Su9-AsCas12a expression but rather a characteristic of the stable cell line obtained. NGS analysis identified mtDNA mutation A8566C with a heteroplasmy level of approximately 40 % (**Fig. 5f**). This mutation results in the amino acid substitutions Gln67His in MT-ATP8 and Ile14Leu in MT-ATP6 (both ATPase genes are overlapping in human mtDNA [4]). We can hypothesize that this mutation may contribute to the decreased respiration observed in the stable cell line; however, further experiments are necessary to characterize this mutation and assess its potential impact on ATP production. Taken together, our findings suggest that temporary expression of the mitoAsCas12a in cultured human cells does not induce mitochondrial dysfunction or mitophagy, which is a crucial consideration for potential applications.

To assess functionality of the mitoAsCas12a, we first investigated mtDNA copy number in the stable cell line treated with crRNAs targeting MT-ND4 and ATP6 (**Fig. 4a, 4b**). Contrary to previous reports showing a decrease in mtDNA copy number following the mitoCas9 treatment [25–29] or even an increase with the mitoLbCas12a [34], we did not observe any significant changes in mtDNA copy number (**Fig. 4e**). We hypothesize that the efficiency of the system is limited by a low level of crRNA import into the mitochondria. So, if mtDNA cleavage did occur, it was not sufficient to cause a measurable shift in mtDNA copy number under our experimental conditions.

However, we successfully detected specific mtDNA cleavages induced by Su9-AsCas12a using more sensitive approaches. Linearized mtDNA, formed as a result of DSBs, was identified through nested LM-PCR (**Fig. 5a-c, Supplementary** Fig. 3c**, 3d**). Additionally, NGS of amplicons confirmed the presence of re-circularized mtDNA molecules carrying the expected deletions of 2.5 and 7 kb (**Fig. 5f, Supplementary** Fig. 3e-g). Along with the detected A8623T substitution (**Fig. 5g, 5h**), which aligns with the natural variability in AsCas12a cleavage position (**Fig. 4a, 4b**), we observed that most of the free ends of mtDNA formed after the cleavage were slightly degraded prior to or during re-ligation (**Fig. 5f, 5g; Supplementary** Fig. 3f**, 3g**). This may possibly be due to the exonuclease activities of PolG and/or MGME1, or to partial double-strand break repair via MMEJ, a process known to introduce mutations, including deletions [7]. Interestingly, mtDNA deletions were detectable even without MGME1 knockdown, indicating that AsCas12a is capable of bypassing mtDNA degradation mechanisms to some extent.

To estimate the efficiency of deletion formation, we compared mtDNA copy number using qPCR with primers targeting regions inside and outside the deletion. However, no difference was observed (**Supplementary** Fig. 4), suggesting that the deletion frequency was low, falling below the detection limit of the applied technique. This result aligns with our previous findings, which showed no change in mtDNA copy number (**Fig. 4e**). Another study reported the presence of InDels in mtDNA after exposure to the mitoCas9 system, with a frequency of less than 0.05 % [28]. Future efforts to improve the efficiency of mtDNA cleavage using the mitoCRISPR system could focus on optimizing crRNA mitochondrial import through chemical modifications that might enhance mitochondrial uptake, as well as employing engineered Cas12a nucleases with improved cleavage activity and specificity [61].

This study represents a significant advance over previous attempts to edit mtDNA using CRISPR-based systems. We demonstrated both mitochondrial import and specific mtDNA cleavage using the mitoAsCas12a system, providing the first evidence of CRISPR-induced deletions in human mtDNA. These findings highlight the potential of mitoAsCas12a for precise and targeted mitochondrial genome editing, opening new avenues for therapeutic interventions in mitochondrial diseases. Future work will focus on refining the specificity and efficiency of the mitoAsCas12a system, thus increasing its potential for clinical and fundamental applications.

## Methods

### Cell culture conditions and mycoplasma screening

HEK-293 cell lines and their derivatives were maintained at 37 °C, 5 % CO_2_, in essential modified Eagle’s medium (EMEM) (Sigma-Aldrich) containing 1 g/L D-glucose, supplemented with 1.5 g/L sodium bicarbonate (Sigma-Aldrich), 0.11 g/L sodium pyruvate (Sigma-Aldrich), 10% fetal bovine serum (Gibco, Fisher Scientific), 100 mg/L Penicillin-Streptomycin (Sigma-Aldrich), and 2.5 mg/L Amphotericin B (Sigma-Aldrich). Prior to each experiment, cell cultures were tested for mycoplasma contamination by PCR with DreamTaq Green PCR Master Mix (Thermo Scientific) (see **Supplementary Table 3** for the list of primers). Total DNA, isolated with NucleoSpin Tissue Mini Kit (Macherey-Nagel), was used as PCR template. Analysis of PCR products was performed by electrophoresis in a 1 % agarose gel (w/v) in 1× TAE buffer.

### Plasmids expressing AsCas12a

All plasmid vectors were designed using SnapGene software V.2.8.3 (GSL Biotech). The plasmid “MitoCas9” [62] was used as a backbone for AsCas12a expression in mammalian cell lines. Cas9 was replaced with the humanized AsCas12a from pY010 (pcDNA3.1-hAsCpf1) (Addgene, #69982) using Gibson Assembly Master Mix (NEB). The vector and the insert were amplified using Phusion High-Fidelity DNA Polymerase (Thermo Scientific) (**Supplementary Table 3**).

The COX8A MTS was exchanged using site-directed mutagenesis. Primers were generated using NEB Base Changer V_1.3.3 (NEB). PCR amplifications were performed using Phusion High-Fidelity DNA Polymerase. The remaining template plasmid was removed by FastDigest *Dpn*I treatment (Thermo Scientific). Ligation was performed using T4 DNA Ligase (Thermo Scientific). A deletion in CMV promoter (Δ5) [37] was introduced by site-directed mutagenesis. ProtParam Expasy online server [63] was used to analyze protein parameters, such as the predicted pI, charge, and grand average of hydropathy (GRAVY) for the amino acid sequences of the MTSs alone and when fused to AsCas12a. For the T-REx-293-Su9-AsCas12a stable cell line generation, the nuclease bearing Su9 MTS was re-cloned into pcDNA5-FRT-TO vector (Invitrogen) using Gibson assembly. Chemically competent *E. coli* XL1-Blue cells were used for plasmids’ replication. Plasmid DNA was isolated using NucleoSpin Plasmid Mini Kit (Macherey-Nagel). The accuracy of the assemblies was verified by Sanger sequencing at Eurofins Genomics.

To express the mitochondrially-targeted AsCas12a, cells (0.8 cm²) were transfected with 100 ng of plasmid DNA encoding the nuclease using Lipofectamine 2000 Transfection Reagent (Invitrogen).

### Western blotting

Cell pellets were resuspended in 1× PBS and sonicated on ice for 10 seconds using a Vibra-Cell VCX 750 Sonicator equipped with a 2 mm probe (Sonics & Materials, Inc.) at a frequency ranging from 20 to 50 kHz. Protein concentration was determined using ROTI Nanoquant (Carl Roth GmbH) with a Bio-Rad SmartSpec Plus UV/Vis Spectrophotometer (Bio-Rad). Denatured protein extracts (20 *μ*g) mixed with Laemmli buffer were analyzed by 10-13 % SDS-PAGE in TGS buffer (25 mM Tris base, 190 mM glycine, 0.1 % SDS, pH 8.3). Following electrophoresis, proteins were transferred onto an Amersham Protran nitrocellulose membrane (GE Healthcare) by semi-dry transfer in 1× transfer buffer (48 mM Tris, 39 mM glycine, 20 % methanol, 0.04 % SDS) using a Trans-Blot Turbo Transfer System (Bio-Rad).

Membranes were blocked using 1× TBST buffer (150 mM NaCl, 50 mM Tris-HCl, 0.1 % Tween-20, pH 7.6) supplemented with 5 % low-fat milk for 1 h at room temperature (RT). Subsequently, membranes were incubated with appropriately diluted primary antibodies in 1× TBST buffer for 1.5 h at RT or overnight at 4 °C, following incubation with diluted HRP-conjugated secondary antibodies for 1 h at RT (refer to **Supplementary Table 4** for the list of antibodies used in the study). Membranes were treated with revelation solution (100 mM Tris-HCl, pH 8.5, 30 % H_2_O_2_, 90 mM coumaric acid, 250 mM luminol), and signals were recorded using a ChemiDoc Touch Imaging System (Bio-Rad). The obtained images were analyzed with Image Lab software (v. 6.0.1) (Bio-Rad).

### Immunofluorescent staining and confocal laser scanning microscopy

Cells (0.15-0.20 cm²) were seeded into 8-well Nunc Lab-Tek slides (Thermo Fisher Scientific) pre-treated with 0.1 % gelatin. After 24 h of incubation, cells were fixed in fixation buffer (4 % paraformaldehyde dissolved in PBS and adjusted to 3 % using EMEM) for 12 minutes at 37 °C. Then, cells were permeabilized with permeabilization buffer (0.5 % (v/v) Triton X-100 in 1× PBS) for 15 minutes at RT. Cells were blocked for 1.5 h in 5 % BSA/PBS solution. Appropriately diluted in blocking solution primary antibodies were added to cells and incubated for 3 h at RT, followed by staining with secondary antibodies conjugated with a fluorescent marker for 1 h at RT (see **Supplementary Table 4** for the list of antibodies used in the study).

Cells were imaged on an LSM700 microscope (Carl Zeiss) using a 63×/1.4 oil objective in 1× PBS. The obtained images were visualized and analyzed using ImageJ 1.52a software [64]. To verify mitochondrial localization of AsCas12a, fluorescence profiles of signals attributed to both mitochondria (TOMM20) and the FLAG-tagged AsCas12a were assessed through a semi-quantitative analysis along a selected 20 *µ*m line using the same software.

### Su9-AsCas12a stable cell line

The Su9-AsCas12a gene was introduced into HEK-293 T-REx cell line nuclear genome using the Flp-In System (Invitrogen). T-REx-293 cell line was transfected in a 6-well plate with a 1:1 ratio of pOG44 Flp-Recombinase expression vector (Invitrogen) and pcDNA5-FRT-TO-Su9-AsCas12a using Lipofectamine 2000 transfection reagent (Invitrogen). Successful integration of the gene was monitored by antibiotic selection with hygromycin B (75 *μ*g/mL, Invitrogen). Cells were allowed to propagate in selective media until cell foci could be identified. Subsequently, individual clones were isolated using the clonal ring anchoring method. The Su9-AsCas12a expression was activated by addition of 100 ng/mL of tetracycline.

### Submitochondrial fractionation of proteins

Crude mitochondria were isolated by differential centrifugation. All procedures were carried out at 4 °C unless specified otherwise. Briefly, 75 cm² of cells were washed once with 1× PBS, then detached and resuspended in 1 mL of prechilled Mito buffer (0.6 M sorbitol, 10 mM HEPES-KOH, pH 7.5, 1 mM EDTA). Cells were disrupted by 30 passages through a 2 mL syringe with a No. 26G needle (0.45×12 mm). Nuclei, cell debris, and unbroken cells were pelleted by centrifugation for 10 min at 1000 g at 4 °C. The procedure was repeated three times, transferring the resulting supernatant each time to a new ice-cold tube. Mitochondria-enriched pellets were obtained after high-speed centrifugation at 15000 g at 4 °C for 30 min.

Submitochondrial fractionation was performed as previously described by **Jeandard et al., 2023** [65] with variations. In brief, mitochondrial pellets were split into three aliquots, each resuspended in Mito buffer, swelling buffer (10 mM HEPES-KOH, pH 7.5, 1 mM EDTA) for mitoplast preparation, and lysis buffer (10 mM HEPES-KOH, pH 7.5, 1 mM EDTA, 0.5 % (w/v) n-dodecyl-*β*-maltoside). Each aliquot was split in two, with one being treated with 50 *μ*g/mL proteinase K (Promega) for 20 min on ice. The reaction was stopped by adding PMSF to a final concentration of 1 mM. To precipitate proteins, ¼ of the total volume of 100 % trichloroacetic acid was added, and samples were incubated on ice for another 10 min. Samples were then centrifuged at 14000 g for 10 min at 4 °C. Supernatants were discarded, and pellets were washed twice with pre-chilled acetone, followed by centrifugation at 14000 g for 10 minutes at 4°C. The resulting supernatants were removed, and protein pellets were dried overnight at RT. Subsequently, protein samples were analyzed by Western blot hybridization.

### AsCas12a crRNAs

crRNAs were synthesized by the team of Dr. Vadim Shmanai, National Academy of Sciences of Belarus, Minsk, Belarus) using an automatic DNA/RNA synthesizer ASM 2000 (Biosset Ltd.) at a 500 nmol scale on 1000 Å Universal Support CPG (Primetech ALC, 41 μmol/g), as described [45]. Additionally, AsCas12a crRNAs were provided by IDT. See **Supplementary Table 5** for the list of crRNAs used in the study.

### *In vitro* cleavage in cell lysate

mtDNA fragments were amplified using Phusion High-Fidelity DNA Polymerase. Genomic DNA isolated from T-REx-293 cell line using NucleoSpin Tissue Mini Kit served as a PCR template (oligonucleotides used for mtDNA amplification are listed in **Supplementary Table 3**).

The expression of the nuclease in the T-REx-293-Su9-AsCas12a cell line was activated at 80 % cells confluency by adding 100 ng/mL tetracycline. After 24 h, cells were washed with 1× PBS and lysed using 50 *µ*L/cm² lysis buffer (20 mM HEPES-KOH pH 7.5, 100 mM KCl, 5 mM MgCl₂, 1 mM DTT, 5 % glycerol, 0.1 % Triton X-100, and Protease Inhibitor Cocktail (Roche Diagnostics)), then rocked for 10 min at 4 °C. To remove cell debris, lysates were centrifuged for 10 min at 1000 g at 4 °C. The resulting supernatants were divided into aliquots and stored at 80 °C.

*In vitro* cleavage reactions were carried out in a total volume of 20 *μ*L and contained 18 *μ*L of the stable cell line lysate, 150 ng of mtDNA fragment, and 50 ng of crRNA (**Supplementary Table 5**). Reactions were incubated for 30 min at 37 °C. After incubation, DNA was purified using NucleoSpin Gel and PCR Clean-up Kit (Macherey-Nagel). Analysis of the cleavage products was performed by electrophoresis in 1 % agarose gel.

### Oxygen consumption analysis

Cells (approximately 10,000) were plated on 8-well Seahorse XF HS Miniplates (Agilent) pretreated with 0.1 % gelatin. 24 h prior to the assay, cells were treated with tetracycline (100 ng/mL) if required. On the day of the assay, culture medium was replaced with Seahorse XF DMEM Medium pH 7.4 (Agilent), supplemented with 1 mM sodium pyruvate, 2 mM glutamine, and 10 mM glucose (pH 7.4), and incubated for 1 hour at 37 °C. The Mito Stress Test Kit (Agilent) was applied using 1.5 *µ*M oligomycin, 2 *µ*M FCCP, and 0.5 *µ*M rotenone/antimycin. Cells were analyzed using the Seahorse XF HS Mini Analyzer (Agilent). After analysis, cells were detached directly in the wells and quantified with the LUNA-II Automated Cell Counter (Logos Biosystems) to normalize the data.

### Inducing deletions in human mtDNA

For MGME1 knockout, the T-REx-293-Su9-AsCas12a cells were transfected in suspension with DsiRNA targeting MGME1 (hs.Ri.MGME1.13.8, IDT) at a final concentration of 40 nM using Lipofectamine RNAiMAX transfection reagent (Invitrogen) in Opti-MEM (Gibco, Fisher Scientific) (see **Supplementary Table 6** for the sequences of siRNA). 6 h after transfection, the transfection medium was replaced with complete medium (EMEM). Three days later, the transfection was repeated under standard adherent cell conditions. 48 h later, the cells were transfected in suspension with a pair of AsCas12a crRNAs targeting human mtDNA (**Supplementary Table 5**) at a final concentration of 125 ng/mL for each crRNA using Lipofectamine 2000 transfection reagent (Invitrogen) in Opti-MEM (Gibco, Fisher Scientific). 6 h after transfection, the transfection medium was replaced by EMEM supplemented with 100 ng/mL of tetracycline to activate the nuclease expression. Two days after, the cells were detached, and total DNA was isolated using the QIAamp DNA Mini Kit (Qiagen).

### qPCR for mtDNA copy number

To quantify mtDNA copy number, qPCR was performed using 20 ng of isolated total DNA with SsoAdvanced Universal SYBR Green Supermix (Bio-Rad) in CFX96 Real-Time System/C1000 Touch Thermal Cycler system (Bio-Rad). The reaction conditions were as follows: an initial denaturation at 95 °C for 3 min, followed by 40 cycles of denaturation at 95 °C for 15 sec and annealing/elongation at 58 °C for 30 sec. For each DNA sample, two separate reactions were prepared, each utilizing primers for either mitochondrial DNA (12S gene) or nuclear DNA (TST gene) amplification (**Supplementary Table 3**). Each qPCR reaction was performed in technical triplicate, and the mean cycle threshold (Ct) value was calculated. qPCR data were analyzed using a Bio-Rad CFX Manager Software (version 3.1; Bio-Rad). Relative mtDNA copy numbers were estimated using the 2^−ΔΔCt^ method.

### Nested LM-PCR

Linker-mediated PCR (LM-PCR) was performed following the procedure described in [9], with modifications. Total DNA (1 *µ*g) was pre-treated with 2 U of Klenow Fragment (Thermo Fisher Scientific) for 15 min at 37 °C, followed by inactivation for 10 minutes at 75 °C. Subsequently, 200 ng of pre-treated DNA was ligated to an asymmetrical double-stranded linker using 400,000 U of T4 DNA ligase (NEB) in a total reaction volume of 60 *µ*L. The ligation reaction was incubated at 16 °C for 16 h, followed by inactivation at 65 °C for 10 min.

To prepare a unidirectional linker, 20 *µ*M primers were mixed in 250 mM Tris-HCl (pH 8). After denaturation at 95 °C for 10 minutes, the temperature gradually lowered. PCR amplification was performed using Phusion High-Fidelity DNA Polymerase. The reaction volume of 12.5 *µ*L contained the following components: 0.25 U of polymerase, 2 mM dNTPs, 0.9 pmol of linker-specific primer, 7.5 pmol of an mtDNA-specific primer (either downstream or upstream of the predicted cleavage site), and 6 ng of linker-ligated total DNA. The amplification protocol was as follows: 98 °C for 30 sec, followed by 42 cycles of 98 °C for 10 sec, 57 °C for 30 sec, and 72 °C for 30 sec, with a final elongation step at 72 °C for 5 min.

PCR products were analyzed by electrophoresis on a 1-2 % agarose gel. Regions of interest from the EtBr-stained gels were excised, and DNA was isolated using NucleoSpin Gel and PCR Clean-up Kit. Elution was performed in 30 *µ*L. 5 *µ*L of the resulting DNA was used for nested LM-PCR with linker-specific primer and mtDNA-specific primer positioned closer to the predicted cleavage site than the previous one, using the same reaction conditions as described above.

### NGS of PCR products

PCR amplifications of mtDNA with primers flanking deletions were performed using Phusion High-Fidelity DNA Polymerase with 100 ng of total DNA as the template (oligonucleotides listed in **Supplementary Table 3**). The amplification protocol was as follows: 98 °C for 30 sec, followed by 34 cycles of 98 °C for 10 sec, 60 °C for 5 sec, and 72 °C for 3 sec. The elongation step was minimized to favorize amplification of mtDNA containing deletions versus WT molecules. PCR products were analyzed in 2% agarose gel. Bands of interest were excised from the gel, and DNA was isolated as described above. Elution was performed in 30 *µ*L. 5 *µ*L of the resulting DNA were used for reamplification with primers containing NGS overhang adapters required for library preparation (#15044223 Rev. B, Illumina), using the same PCR program, including a final elongation step at 72 °C for 5 min. DNA purification was performed with NucleoMag kit for clean-up and size selection using NucleoMag SEP Mini separation plate (Macherey-Nagel). 4 ng of isolated DNA were used to complete the libraries by attaching indexes with xGen NXT UDI Primers (IDT), following an amplification protocol of 98 °C for 30 sec, followed by 8 cycles of 98 °C for 10 sec, 72 °C for 30 sec, and a final elongation step at 72ss°C for 5 min. The reactions were again purified with magnetic beads, and the resulting DNA was analyzed using 2100 Bioanalyzer (Agilent). 150-bp paired-end reads with a coverage of no less than 3000 were generated using MiSeq (Illumina) at Gene expression analysis platform (IBMP, Strasbourg).

The obtained reads were analyzed using Galaxy server (version 23.1.rc1) (GCC2022, USA). Adapter sequences were trimmed, and nucleotides with a quality score of less than 20 were filtered out using Trimmomatic (version 0.38) [66]. Reads were then mapped to the reference mtDNA genome (GenBank NC_012920 gi:251831106) or to variants bearing deletions 8624-10928 or 8624-15738 using BWA-MEM2 (version 2.2.1) [67]. The resulting alignments were visualized using IGV (version 2.16.1) [68]. Sequencing data were processed using BAMtools (version 2.5.1) [69] and SAMtools (version 1.13) [70].

## Data availability

All data presented in the manuscript are available upon request. NGS data have been deposited in the GitHub repository under accession link https://github.com/NataliaNikitchina/NGS_data_Nikitchina_et_al_2024.

## Acknowledgements

The authors acknowledge Egor Ulashchik and Vadim Shmanai (National Academy of Sciences of Belarus) for the chemical synthesis of AsCas12a crRNAs. We are grateful to Anna Smirnova (UMR7156 GMGM, CNRS/University of Strasbourg) for technical assistance in fluorescent microscopy and to Sandrine Koechler (IBMP, CNRS/University of Strasbourg) for performing NGS. This work was supported by the Interdisciplinary Thematic Institute (ITI) Integrative Molecular and Cellular Biology (IMCBio+), as part of the ITI 2021–2028 program of the University of Strasbourg, CNRS, Inserm and IdEx Unistra (ANR-10-IDEX-0002). I.M. was supported by grant from Russian Science Foundation No. 23-45-10010.

## Author contributions

N.N, A.M.H. conducted the experiments, N.S. contributed to cell culture and stable cell line generation, N.N. performed visualization, writing, drafting, and I.M, N.E. and I.T. were responsible for conceptualization, data analyses, writing, review and editing.

## Competing interests

The authors declare no competing interests.

## Notes

https://github.com/NataliaNikitchina/NGS_data_Nikitchina_et_al_2024

### Competing Interest Statement

The authors have declared no competing interest.

