## Supplementary Table, Supplementary Fig. for "Targeted deletions in human mitochondrial DNA engineered by Type V CRISPR-Cas12a system"

Nikitchina *et al.*, 2024

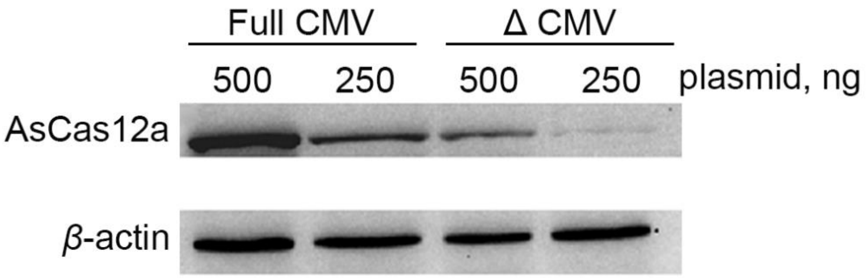


**Supplementary Figure 1** AsCas12a levels decrease when expressed from a plasmid with a weakened CMV promoter. Western blot analysis of T-REx-293 cells 24h after transfection with COX8A-3xFLAG-AsCas12a plasmids, bearing either the full-sized or weakened (Δ5) CMV promoter as indicated above the panel. The amount of plasmid used for the transfection is indicated in ng. AsCas12a was detected using FLAG-tag-specific antibodies, and *β*-actin served as a loading control.

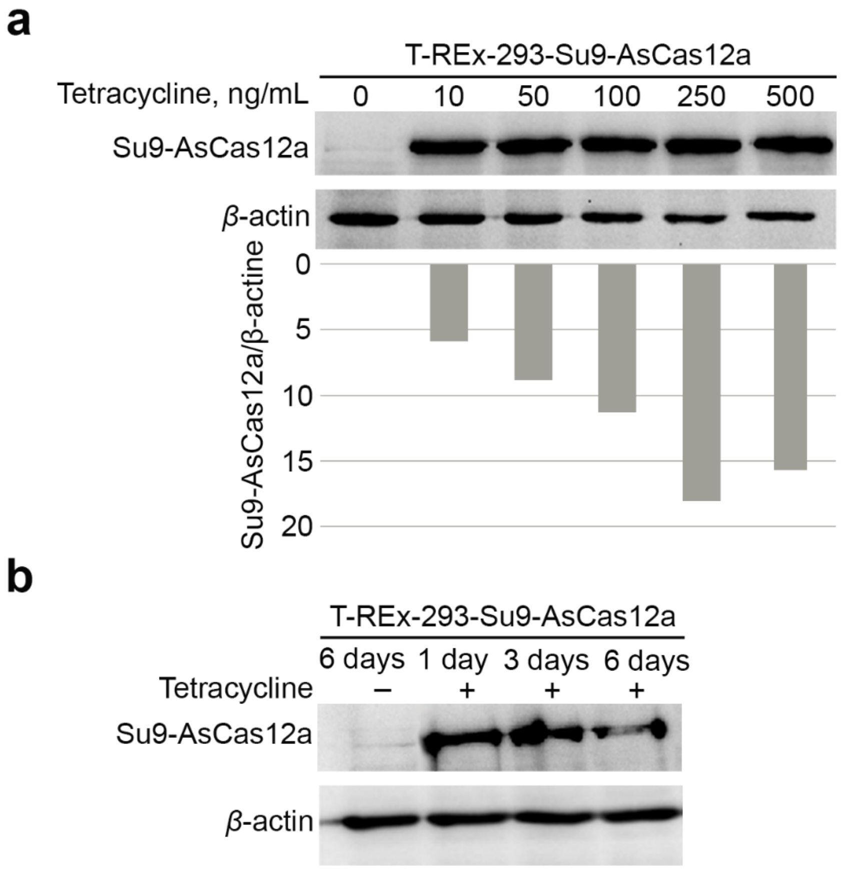


**Supplementary Figure 2** Analysis of Su9-AsCas12a expression in the stable cell line. **a** Western blot analysis performed on total cell lysates of the T-REx-293-Su9-AsCas12a stable cell line, induced with various tetracycline concentrations (as indicated above the panel). Relative expression of Su9-AsCas12a normalized by actin is present on the panel below. Su9-AsCas12a was detected using FLAG-tag-specific antibodies, and *β*-actin served as a loading control. n=1. **b** Similar to (**a**), but cells were detached at different time points after activating Su9-AsCas12a expression to assess its stability. The nuclease expression was induced by adding 100 ng/mL of tetracycline. n=1.

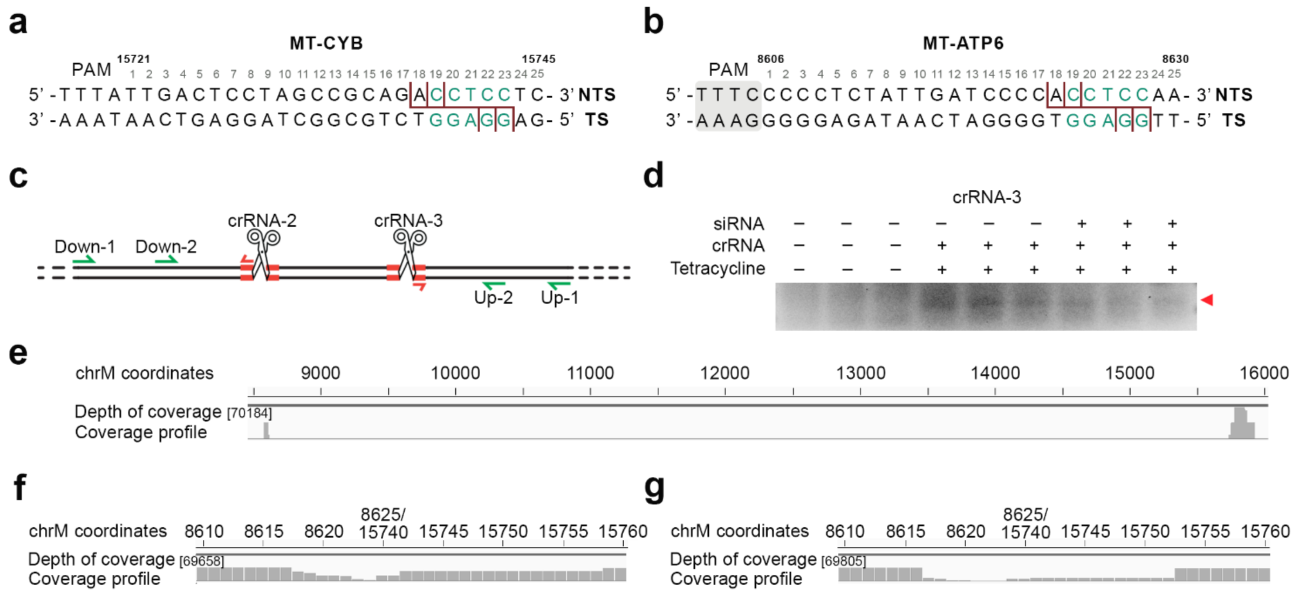


**Supplementary Figure 3** Inducing deletion in human mtDNA by using Su9-AsCas12a in complex with crRNAs targeting MT-CYB and MT-ATP6. **a**, **b** Representation of the cleavage sites in human mtDNA that can be induced by AsCas12a in complex with crRNA-3 targeting MT-CYB (**a**), and crRNA-2 targeting MT-ATP-6 (**b**), designed to leave complementary ends (corresponding nucleotides are highlighted in blue). Possible cleavage sites on both DNA strands are indicated with red lines. The corresponding nucleotide positions in WT human mtDNA are shown above. **c** Schematic representation of nested LM-PCR, where linearized mtDNA molecules are ligated with an asymmetric double-stranded linker to identify cleavage sites. Nested LM-PCR involves two rounds of PCR using primers specific to both the linker (red arrows) and the mtDNA regions either downstream or upstream of the break sites (green arrows). **d** Nested LM-PCR detected the cleavage generated by Su9-AsCas12a in complex with crRNA-3 in living cells. Transfections with crRNA and siRNA to MGME1, as well as the nuclease expression activation, are indicated above the gel. **e** Example of the coverage profile for the sample containing Su9-AsCas12a and crRNAs 2 and 3 with MGME1 downregulation, obtained through short-read NGS of a PCR amplicon and visualized using IGV. Alignment was done to WT human mtDNA. The depth of coverage and chrM coordinates are indicated. **f**, **g** Similar to (**e**), but alignment was performed against an mtDNA reference bearing the 8624-15738 bp deletion for samples treated with Su9-AsCas12a and crRNAs 2 and 3, either without (**f**) or with (**g**) MGME1 downregulation via siRNA, to evaluate the deletion boundaries. Representation of n = 2 for independent biological replicates.

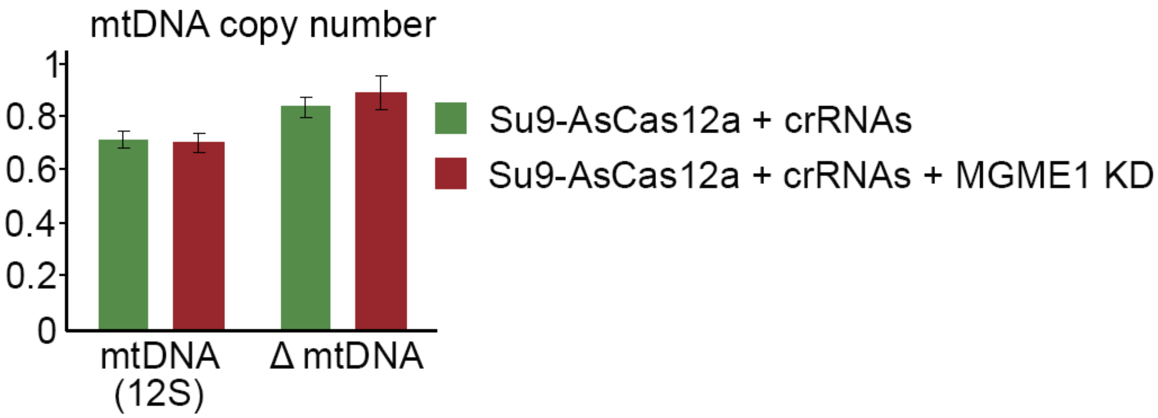


**Supplementary Figure 4** Estimation of deletion formation efficiency by AsCas12a and crRNAs targeting MT-ND4 and NMT-ATP6. qPCR analysis of mtDNA copy number in the T-REx-293-Su9-AsCas12a stable cell line ("Su9-AsCas12a") with or without MGME1 knockdown ("MGME1 KD"), conducted 48 h after treatment with Su9-AsCas12a and crRNAs 1 and 2 ("crRNAs") with primers targeting mtDNA regions inside and outside of the deletion region. Relative mtDNA copy number was quantified using the $2^{-\Delta\Delta Ct}$ method, with MOCK transfection of the stable cell line without MGME1 downregulation used for normalization (mtDNA copy number = 1). Error bars represent the mean ± s.d. of n = 3 for technical replicates.

**Supplementary Table 1**

**Mitochondrial targeting signals (MTSs) used in the study**

| MTS name | Amino acid sequence | Length |
|---|---|---|
| SOD2 | MLSRAVCGTSRQLAPVLGYLGSRQ | 24 |
| COX8A | MSVLTPLLLRGLTGSARRLPVPRAKIHSL | 29 |
| ATG4D | MKFKAKFLTAWNNVKYGWVVKSRTSFSKISSIHLCGRRYRF | 41 |
| Su9 | MASTRVLASRLASRMAASAKVARPAVRVAQVSKRTIQTGSPLQTLKRTQMTSIVNATTRQAFQKRAYSS | 69 |

**Supplementary Table 2**

**Characterization of the AsCas12a nuclease fused to various MTSs**

| Construct | MTS charge[a] | MTS pI[b] | MTS GRAVY[c] | Full Charge[d] | Full pI[e] | Full GRAVY[f] |
|---|---|---|---|---|---|---|
| SOD2-3xFLAG-AsCas12a | +5 | 10.76 | 0.163 | -1 | 8.01 | -0.492 |
| COX8A-3xFLAG-AsCas12a | +3 | 12.48 | 0.362 | +1 | 7.65 | -0.507 |
| ATG4D-3xFLAG-AsCas12a | +10 | 11.17 | -0.337 | +6 | 8.31 | -0.520 |
| Su9-3xFLAG-AsCas12a | +13 | 12.55 | -0.272 | +9 | 8.53 | -0.513 |

[a] Total number of positively charged residues, excluding the negative ones, for the MTS alone.
[b] Theoretical isoelectric point of the MTS alone.
[c] Grand average of hydropathicity of the MTS alone.
[d] Total number of positively charged residues, excluding the negative ones, for the full construct.
[e] Theoretical isoelectric point of the full constructs.
[f] Grand average of hydropathicity of the full constructs.

**Supplementary Table 3**

**DNA oligonucleotides used in the study**

| Primer name | Sequence: 5’- 3’ | Annealing temperature, °C |
|---|---|---|
| *Gibson Assembly* | | |
| F_pY010 | TGACGATGACAAGCTCATGACACAGTTCGAGGGC | 60 |
| R_ pY010 | CCTGTCAGCCCTGCTGTTGCGCAGCTCCTGGAT | 60 |
| F_ pMito-Cas9 | AGCAGGGCTGACAGGTCC | 60 |
| R_ pMito-Cas9 | GAGCTTGTCATCGTCATCCTTG | 60 |
| F_FRT_As | GCGCAACTAACCCGCTGATCAGCCTCG | 60 |
| R_FRT_As | GGAGGCCATACTAAACGAGCTCGTCGACGA | 60 |
| F_SuAsCas12ato FRT | CGTTTAGTATGGCCTCCACTCGTGT | 60 |
| R_SuAsCas12ato FRT | TCAGCGGGTTAGTTGCGCAGCTCCTGGA | 60 |
| *Site-directed mutagenesis* | | |
| F_(d5)CMV | TGTCGTAACAACTCCGCCCCATTG | 60 |
| R_(d5)CMV | CGGAACTCCATATATGGGCTATG | 60 |
| F_As_SOD2 | CTGGCTCCGGTTTTGGGGTATCTGGGCTCCAGGCAG GACTACAAAGACCATGACGG | 64 |
| R_As_SOD2 | CTGCCTGCTGGTGCCGCACACTGCCCGGCTCAACAT GCTAGCGGATCTGACGGT | 64 |
| F_As_ATG4D | AAGCCGGACCAGCTTTAGCAAGATCTCCAGCATCCA CCTCTGTGGCCGCCGCTACCGTTTCGACTACAAAGA CCATGACGG | 64 |
| R_As_ATG4D | TTAACCACCCAACCGTACTTGACGTTGTTCCAGGCTG TCAGGAACTTGGCCTTGAACTTCATGCTAGCGGATCT GACGGT | 64 |
| F_As_Su9 | ATCCAGACTGGCTCCCCCCTCCAGACCCTCAAGCGC ACCCAGATGACCTCCATCGTCAACGCCACCACCCGC | 64 |
| R_As_Su9 | GGTGCGCTTGCTGACCTGAGCAACGCGGACAGCAG GGCGGGCAACCTTGGCGGAAGCAGCCATCCGGGAG GCCAGGCGAGAGGCGAGGACACGAGTGGAGGCCAT GCTAGCGGATCTGAC | 64 |

continued

| Primer name | Sequence: 5’- 3’ | Annealing temperature, °C |
| --- | --- | --- |
| *Mycoplasma screening* | | |
| RNA5 | AGAGTTTGATCCTGGCTCAGGA | 61 |
| RNA3 | ACGAGCTGACGACAACCATGCAC | 61 |
| UNI – | TAATCCTGTTTGCTCCCCAC | 61 |
| *In vitro cleavage tests* | | |
| L7831 | CATCCTTTACATAACAGACG | 58 |
| H13555 | AGGCGTTTGTGTATGATATGTTTG | 58 |
| *qPCR for mtDNA copy number* | | |
| F_12S | GCACTTAAACACATCTCTGCC | 58 |
| R_12S | TGAGATTAGTAGTATGGGAGTGG | 58 |
| F_TST | GACTGGACTCGGGCCATATC | 58 |
| R_TST | ACGTGGCAATGAGAGGCTG | 58 |
| L8542 | TTCGCTTCATTCATTGCCCCC | 58 |
| H11024 | GTGGTTCACTGGATAAGTGGCG | 58 |
| *Nested LM-PCR* | | |
| LMPCR1 | GCGGTGACCCGGGAGATCTGTATTC | 57 |
| LMPCR2 | GAATACAGATC | 57 |
| L7831 | CATCCTTTACATAACAGACG | 57 |
| L8342 | GAACCAACACCTCTTTACAGTG | 57 |
| H11870 | GGGGTAAGGCGAGGTTAGC | 57 |
| H11381 | AAGTGGAGTCCGTAAAGAGG | 57 |
| H16000 | CTTAGCTTTGGGTGCTAATGG | 57 |
| H15929 | TCCGGTTTACAAGACTGGTGTA | 57 |

continued

| Primer name | Sequence: 5’- 3’ | Annealing temperature, °C |
|---|---|---|
| *PCR for NGS* | | |
| L8542 | TTCGCTTCATTCATTGCCCCC | 60 |
| H11015 | TGGATAAGTGGCGTTGGCTTG | 60 |
| L8542-adptr | TCGTCGGCAGCGTCAGATGTGTATAAGAGACAGTTCGCTTCATTCATTGCCCCC | 72 |
| H11015-adptr | GTCTCGTGGGCTCGGAGATGTGTATAAGAGACAGTGGATAAGTGGCGTTGGCTTG | 72 |
| L8581 | CGCAGTACTGATCATTCTATTTCC | 60 |
| H15929 | TCCGGTTTACAAGACTGGTGTA | 60 |
| L8581-adptr | TCGTCGGCAGCGTCAGATGTGTATAAGAGACAGCGCAGTACTGATCATTCTATTTCC | 72 |
| H15929-adptr | GTCTCGTGGGCTCGGAGATGTGTATAAGAGACAGTCCGGTTTACAAGACTGGTGTA | 72 |

**Supplementary Table 4**

**Antibodies used in the study**

| Antibodies name | Provider | Reference |
|---|---|---|
| *Primary antibodies* | | |
| Mouse monoclonal anti-FLAG (clone M2) | Sigma-Aldrich | F1804 |
| Goat polyclonal anti-Actin (C-11) | Santa Cruz Biotechnology | sc-1615 |
| Rabbit polyclonal anti-TOMM20 (FL-145) | Santa Cruz Biotechnology | sc-11415 |
| Rabbit polyclonal anti- PNPT1/PNPase | Abcam | ab96176 |
| Rabbit polyclonal anti-OPA1 | Proteintech Group | 27733-1-AP |
| Mouse monoclonal anti-AsCas12a | Sigma-Aldrich | MABE1823 |
| Mouse monoclonal anti-COX1 (clone 1D6E1A8) | Abcam | ab14705 |
| Mouse monoclonal anti-VDAC1 (clone B-6) | Santa Cruz Biotechnology | sc-390996RR |
| Mouse monoclonal anti-BECN1/Beclin-1 (E-8) | Santa Cruz Biotechnology | sc-48341 |
| Mouse monoclonal anti-PINK1 (C-3) | Santa Cruz Biotechnology | sc-518052 |
| Rabbit polyclonal anti-MGME1 | Thermo Fisher Scientific | 23178-1-AP |
| *Secondary antibodies* | | |
| Rabbit polyclonal anti-Goat IgG HRP | Sigma-Aldrich | A8919 |
| Rabbit polyclonal anti-Mouse IgG HRP | Agilent | P0260 |
| Donkey polyclonal anti-Rabbit IgG HRP | Sigma-Aldrich | NA934V |
| Goat polyclonal anti-Mouse IgG-Alexa Fluor 488 | Thermo Fisher Scientific | A-11001 |
| Goat polyclonal anti-Rabbit IgG-Alexa Fluor 647 | Thermo Fisher Scientific | A-21246 |

**Supplementary Table 5**

**crRNAs used in the study**

| crRNA name | Sequence: 5’- 3’ | Length, bp |
|---|---|---|
| crRNA-1 | UAAUUUCUACUCUUGUAGAUGCUGUUCCCCAACCUUUUCC | 40 |
| crRNA-2 | UAAUUUCUACUCUUGUAGAUCCCCUCUAUUGAUCCCCACC | 40 |
| crRNA-2 + 3 | UAAUUUCUACUCUUGUAGAUCCCCUCUAUUGAUCCCCACCUCC | 43 |
| crRNA-3 + 3 | UAAUUUCUACUCUUGUAGAUUUGACUCCUAGCCGCAGACCUCC | 43 |

**Supplementary Table 6**

**Knock-down of genes by siRNA**

| Gene | siRNA duplex sequence: 5’- 3’ |
|---|---|
| MGME1 | rArGrCrCrUrUrGrGrArArArGrCrArUrArCrUrUrUrCrArCCC<br>rGrGrGrUrGrArArArGrUrArUrGrCrUrUrUrCrCrArArGrGrCrUrUrC |
